# Stimulus set meaningfulness and neurophysiological differentiation: a functional magnetic resonance imaging study

**DOI:** 10.1101/013078

**Authors:** Melanie Boly, Shuntaro Sasai, Olivia Gosseries, Masafumi Oizumi, Adenauer Casali, Marcello Massimini, Giulio Tononi

## Abstract

A meaningful set of stimuli, such as a sequence of frames from a movie, triggers a set of different experiences. By contrast, a meaningless set of stimuli, such as a sequence of ‘TV noise’ frames, triggers always the same experience – of seeing ‘TV noise’ – even though the stimuli themselves are as different from each other as the movie frames. We reasoned that the differentiation of cortical responses underlying the subject’s experiences, as measured by Lempel-Ziv complexity (incompressibility) of functional MRI images, should reflect the overall meaningfulness of a set of stimuli for the subject, rather than differences among the stimuli. We tested this hypothesis by quantifying the differentiation of brain activity patterns in response to a movie sequence, to the same movie scrambled in time, and to ‘TV noise’, where the pixels from each movie frame were scrambled in space. While overall cortical activation was strong and widespread in all conditions, the differentiation (Lempel-Ziv complexity) of brain activation patterns was correlated with the meaningfulness of the stimulus set, being highest in the movie condition, intermediate in the scrambled movie condition, and minimal for ‘TV noise’. Stimulus set meaningfulness was also associated with higher information integration among cortical regions. These results suggest that the differentiation of neural responses can be used to assess the meaningfulness of a given set of stimuli for a given subject, without the need to identify the features and categories that are relevant to the subject, nor the precise location of selective neural responses.

## Introduction

When one watches a movie on a TV screen, as the movie frames flow by, one sees a succession of different scenes, each containing a different assortment of meaningful objects and events. It is fair to assume that the different scenes one experiences (phenomenological differentiation) are due to different patterns of activity in relevant parts of the brain (neurophysiological differentiation), including regions that respond to faces, places, story lines, and so on. Ultimately, of course, such different patterns of activity are related to differences among the physical stimuli to which the brain is exposed (stimulus set differentiation) – in this case, the movie frames.

If instead one watches for a while a TV out of tune, as the black and white pixels flicker in front of the eyes, one sees all the time the same flickering image of ‘TV noise.’ As with the movie, each ‘TV noise’ frame is made up of a different configuration of black and white pixels (stimulus set differentiation), each of which will trigger different patterns of activity in the retina and possibly elsewhere in the visual system. However, it is fair to assume that the parts of the brain that are relevant for what one sees consciously will respond in the same way to different noise patterns (lack of neurophysiological differentiation), since what one sees and its meaning stay the same (TV noise, lack of phenomenological differentiation).

By comparing the differentiation of brain responses triggered by a movie with those triggered by its pixel-scrambled version, one might therefore obtain a neurophysiological index of how much more subjectively meaningful the movie is, compared to the TV noise, even though the objective differences in the set of stimuli are comparable [1]. Importantly, measuring the increase in neurophysiological differentiation to a movie compared to ‘TV noise’ in an individual subject does not require knowing which brain region responds to which movie features, nor does it require that different subjects respond in similar ways to the same features - it only requires that they respond in a more differentiated manner to a set of meaningful stimuli than to a less meaningful one.

## Material and Methods

### Subjects and Ethics Statement

Six healthy participants (N=6; 4 females; age range, 28-48 years) from the University of Wisconsin–Madison community participated in the study. All subjects provided informed consent following the procedures approved by the Health Sciences Institutional Review Board of the University of Wisconsin–Madison. Subjects had normal or corrected-to-normal vision, no contraindications for MRI, and no reported neurological or psychiatric history.

### Stimuli

Stimuli were compiled from a classic silent film [Charlie Chaplin’s City Lights (1931)] [2]. We used a silent film in order to avoid potential complications associated with temporal scrambling of sound [2]. The clips were of 256 seconds (~4 min) duration and resampled using Windows Movie Maker at a rate of 30 Hz. For the scrambled movie condition, the 4 min film sequence was subdivided into segments and scrambled in time at a 4 sec time scale. The original film was first divided into 4 seconds segments then randomly resampled using a Matlab custom script. For the ‘TV noise’ condition, movies were spatially scrambled by randomly re-assigning each individual image pixels location in space using a Matlab custom script. This procedure was meant to preserve the first-order statistics of the stimuli. The temporal sequence and sampling rate of the ‘TV noise’ frames was also kept the same as in the original movie.

A general issue in experiments comparing scrambled to unscrambled images is that higher order image statistics are usually not fully matched even when using conservative procedures such as Fourier scrambling [3]. Ideally, multiple low-level scrambling procedures could be employed and compared, but this was not feasible in this initial study employing 30 repeated trials per condition. Therefore, in the present experiment we employed random spatial scrambling as the lowest-level fully structure less baseline [1] against which to compare to movie and scrambled movie data. ‘TV noise’ represents the lowest-level baseline because ‘TV noise’ does in fact look always the same. This is not true for Fourier and other scrambling procedures that preserve low-level correlations: Fourier scrambled frames look clearly different, although they do not trigger any discernible high-level category such as contours or objects. On the other hand, ‘TV noise’ can certainly reach far enough into the brain to trigger the appropriate experience (of ‘TV noise’), while presumably producing differentiated patterns of activity at least in the retina. Only ‘TV noise’, then, truly removes all spatial and temporal correlations to which the brain is attuned and, having no structure, it has no differentiated meanings for the brain to pick up – whether low-or high-level.

The block design itself was made of 10 presentations of 20 sec. of the original movie, 20 sec. of a 4 sec. time-scrambled version of this 20 sec. movie sequence, and 20 sec. of the ‘TV noise’ corresponding to the 20 sec. movie, in counterbalanced order. Each 20 sec. of stimulus presentation was preceded and followed by 10 sec. of black screen.

### Experimental paradigm

While in the fMRI scanner, subjects viewed thirty presentations of 4 min segments of movie, scrambled movie, and ‘TV noise’ in counterbalanced order. Subjects watched the movie twice before starting the experiment to become familiar with its content. The experiment took place in 2 separate days (except for one subject, where it was spread over 3 afternoons). Each subject also underwent the block design session and a 4 min resting state scanning session. During the whole experiment, subjects were instructed to focus on the visual stimuli and avoid mind wandering. Online vigilance monitoring was performed using eye tracking and simultaneous electroencephalographic recordings using a 32-electrode Brain Amp magnetic resonance compatible EEG setup. These data are not reported in the present article, but were used to ensure that subjects had remained vigilant and focused throughout the experiment.

### fMRI data acquisition

Functional MRI time series were acquired using a 3 Tesla GE MR scanner. Multislice T2*-weighted fMRI images (TR 1100 ms, TE 14 ms, 29 slices, with a slice thickness 4 mm and an inter-slice gap of 1 mm) and a structural T1-weighted sequence were acquired in each subject. A single block design consisted in 828 scans. Thirty times 235 scans were acquired for repeated stimulus presentations of movie, scrambled movie and ‘TV noise’ conditions (for a total of ninety times 4 min sessions i.e. seven hours of scanning per subject). A high-resolution T1 image was also acquired in each volunteer at the end of the whole experiment for coregistration to the functional data. During data acquisition, subjects wore earplugs and headphones and were in a comfortable supine position.

### Block design analysis

Figure 1, Top Panel, displays the experimental paradigm used for block design analysis. In this paradigm, 20 sec sequences of movie, scrambled movie, or ‘TV noise’ were presented in alternation with a black screen baseline. The block design analysis was expected to show significant increases in the mean activity of many voxels for each of the three stimulus sequences compared to the black screen baseline.

**Figure 1.**
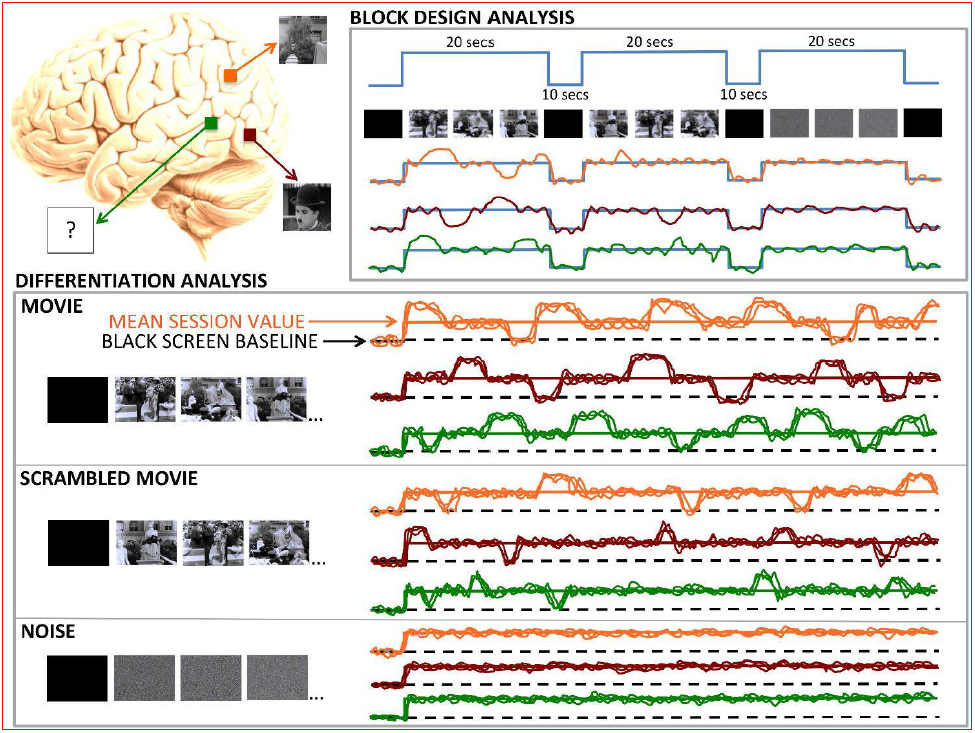
Experimental paradigm. Left panel: Schematic representation of a human brain with 3 representative voxels whose fMRI BOLD activity will be measured in a block design analysis and in a differentiation analysis. The red voxel is known to respond to faces, the orange voxel to places, and the green voxel has unknown selectivity. The colored traces in the right side panels represent the expected BOLD signal of these representative voxels during the fMRI experiments. Right, Top Panel: in the block design paradigm, 20 seconds sequences of movie, scrambled movie, or ‘TV noise’ are presented in alternation with a black screen baseline. The block design analysis is expected to reveal significant increases in the mean activity of the three pictured voxels for each of the three stimulus sequences compared to the black screen baseline. Bottom Panel: in the differentiation analysis paradigm, a 4 min sequence of movie, scrambled movie or ‘TV noise’ is presented to the subjects, each sequence repeated 30 times across different scanning sessions (only 3 of these repetitions are depicted, corresponding to 3 BOLD activity traces per voxel). In all three conditions, we expect an overall activation with respect to the black screen baseline similar to that in the block design paradigm. However, unlike the block design analysis, the differentiation analysis focuses on systematic time-locked increases or decreases in activity with respect to: each voxel’s the black screen baseline (dashed black line); ii) each voxel’s mean activity during the 4 min sequence. Movie sequence: for each voxel, we expect systematic time-locked increases and decreases of activity across the session (neurophysiological differentiation over time); moreover, we expect different voxels to show different patterns of systematic activations/deactivations in response to different movie frames (neurophysiological differentiation over space). Altogether, high neurophysiological differentiation in space and time (many different spatio-temporal patterns) is expected to go along high phenomenological differentiation (many different experiences). Scrambled movie sequence: we expect intermediate levels of neurophysiological differentiation, corresponding to intermediate levels of phenomenological differentiation. TV noise sequence: we expect no or minimal systematic time-locked incease or decreases in activity. Low neurophysiological differentiation (a single, unchanging pattern of activation/deactivation) corresponds to low phenomenological differentiation (a single, unchanging experience of ‘TV noise’). Spontaneous fluctuations in BOLD activity from scan to scan are also expected in the ‘TV noise’ session, but they will not be time locked to specific ‘TV noise’ frames, which cortical regions treat as equivalent.

For this analysis, fMRI data were analyzed using Statistical Parametric Mapping (SPM, www.fil.ion.ucl.ac.uk/spm). Spatial preprocessing of functional scans included realignment, normalization to an MNI template, and smoothing using an 8 mm FWHM Gaussian kernel. After preprocessing, the onsets of movie, scramble movie and noise presentation were modeled using a box-car design and convolved with a canonical hemodynamic response function. In each subject, we then performed a T-test searching for differences in activation between the movie, scrambled movie and ‘TV noise’ sequences as compared to the black screen baseline. Individual results were thresholded at whole-brain FWE p<0.05 (see representative subject in Figure 2). Each subject’s unthresholded T contrast images for the movie, scrambled movie and ‘TV noise’ were also entered in a second level random effects group analysis, using a full factorial design with non-sphericity correction. Group results were computed T contrasts and thresholded at whole-brain FWE p<0.05 (Table 1).

**Table 1.** Block design overall activation versus a black screen baseline

|  | Side | Area | BA | X | Y | Z | Z value | P value |
| --- | --- | --- | --- | --- | --- | --- | --- | --- |
| Movie<br>907 significant voxels |  |  |  |  |  |  |  |  |
|  | R | MOG | 37 | 46 | -72 | 0 | 6.47 | <0.001 |
|  |  | MOG | 19 | 42 | -80 | 0 | 5.83 | <0.001 |
|  | L | MTG | 39 | -44 | -70 | 8 | 5.77 | <0.001 |
|  |  | MOG | 19 | -46 | -80 | 12 | 5.75 | <0.001 |
|  |  | MOG | 37 | -50 | -74 | 2 | 5.63 | <0.001 |
|  | L | MOG | 18 | -34 | -90 | 8 | 5.60 | 0.001 |
|  |  | Cuneus | 18 | -12 | -102 | 4 | 5.33 | 0.002 |
|  |  | Cuneus | 18 | -18 | -102 | 6 | 5.32 | 0.002 |
|  | L | Precentral Gyrus | 6 | -58 | 2 | 30 | 5.19 | 0.004 |
|  | R | Lingual gyrus | 17 | 14 | -94 | -10 | 5.13 | 0.005 |
|  | L | Fusiform gyrus | 20 | -42 | -38 | -20 | 5.08 | 0.007 |
|  | R | Fusiform Gyrus | 37 | 36 | -52 | -14 | 5.03 | 0.009 |
|  | R | Fusiform Gyrus | 19 | 28 | -70 | -10 | 4.98 | 0.010 |
|  | L | Insula | 13 | -38 | -2 | -10 | 4.80 | 0.022 |
|  | R | Postcentral Gyrus | 2 | 34 | -32 | 40 | 4.73 | 0.028 |
|  | R | Postcentral Gyrus | 2 | 40 | -30 | 44 | 4.72 | 0.030 |
|  | R | Cuneus | 18 | 18 | -100 | 8 | 4.58 | 0.050 |
| Scrambled<br>movie<br>947 significant voxels |  |  |  |  |  |  |  |  |
|  | R | MOG | 37 | 46 | -72 | 0 | 6.59 | <0.001 |
|  |  | MOG | 19 | 42 | -80 | 0 | 5.78 | <0.001 |
|  |  | MOG | 18 | 36 | -90 | 0 | 5.77 | <0.001 |
|  | L | MTG | 39 | -44 | -70 | 8 | 5.87 | <0.001 |
|  |  | MOG | 19 | -46 | -80 | 12 | 5.87 | <0.001 |
|  |  | MOG | 37 | -50 | -74 | 2 | 5.76 | <0.001 |
|  | L | MOG | 19 | -30 | -94 | 4 | 5.56 | 0.001 |
|  |  | Cuneus | 18 | -12 | -102 | 4 | 5.41 | 0.002 |
|  |  | Cuneus | 17 | -10 | -100 | -4 | 5.36 | 0.002 |
|  | R | Lingual Gyrus | 17 | 12 | -94 | -10 | 5.41 | 0.002 |
|  | R | Fusiform Gyrus | 37 | 36 | -52 | -14 | 4.91 | 0.014 |
|  | R | Fusiform Gyrus | 19 | 28 | -70 | -10 | 4.86 | 0.017 |
|  | R | Cuneus | 18 | 18 | -100 | 8 | 4.70 | 0.032 |
|  | L | MOG | 18 | -28 | -82 | -12 | 4.65 | 0.040 |
|  | L | Fusiform Gyrus | 37 | -42 | -40 | -20 | 4.61 | 0.045 |
|  | L | MOG | 18 | -24 | -86 | -14 | 4.60 | 0.047 |
| 'TV Noise'<br>104 significant voxels |  |  |  |  |  |  |  |  |
|  | R | Cuneus | 18 | -18 | -102 | -8 | 5.69 | <0.001 |
|  | L | Cuneus | 18 | -10 | -102 | -4 | 5.49 | 0.001 |
|  | R | Lingual gyrus | 17 | 14 | -94 | -10 | 5.36 | 0.002 |
|  | R | Cuneus | 18 | 24 | -98 | -2 | 5.34 | 0.002 |
|  | L | Lingual Gyrus | 17 | -6 | -90 | -10 | 4.62 | 0.043 |
| Movie vs 'TV Noise'<br>327 significant voxels |  |  |  |  |  |  |  |  |
|  | R | MOG | 37 | 46 | -72 | 0 | 5.58 | 0.001 |
|  |  | MOG | 19 | 42 | -80 | 0 | 5.46 | 0.001 |
|  | L | MOG | 18 | -34 | -90 | 8 | 5.42 | 0.002 |
|  | L | Fusiform Gyrus | 19 | -40 | -70 | -14 | 5.26 | 0.003 |
|  |  | MTG | 37 | -46 | -66 | 6 | 5.14 | 0.005 |
|  |  | MOG | 37 | -50 | -74 | 0 | 5.10 | 0.006 |
|  | L | MOG | 19 | -46 | -80 | 12 | 5.23 | 0.004 |
|  | L | Precuneus | 31 | -24 | -76 | 26 | 4.88 | 0.016 |
|  | L | Fusiform Gyrus | 20 | -42 | -34 | -22 | 4.86 | 0.017 |
|  |  | Fusiform Gyrus | 37 | -40 | -42 | -22 | 4.79 | 0.023 |
|  | R | Fusiform Gyrus | 37 | 36 | -52 | -14 | 4.74 | 0.027 |
|  | R | Cingulate Gyrus | 31 | 8 | -50 | 42 | 4.68 | 0.035 |
|  | R | Precuneus | 7 | 22 | -68 | 32 | 4.64 | 0.041 |
| Scrambled movie vs 'TV Noise'<br>322 significant voxels |  |  |  |  |  |  |  |  |
|  | R | MOG | 37 | 46 | -72 | 0 | 5.68 | <0.001 |
|  |  | MOG | 19 | 42 | -80 | 0 | 5.39 | 0.002 |
|  | L | Fusiform Gyrus | 19 | -40 | -70 | -14 | 5.32 | 0.002 |
|  |  | MOG | 19 | -46 | -80 | 12 | 5.31 | 0.003 |
|  |  | MOG | 37 | -50 | -74 | 0 | 5.18 | 0.005 |
|  | L | MOG | 18 | -34 | -90 | 8 | 5.32 | 0.002 |
|  | L | Precuneus | 31 | -24 | -74 | 28 | 4.94 | 0.012 |
|  | R | Fusiform Gyrus | 37 | 36 | -52 | -14 | 4.63 | 0.043 |
| Movie vs Scrambled movie<br>0 significant voxel |  |  |  |  |  |  |  |  |

Block design overall activation group results for movie, scrambled movie and 'TV noise' as compared to a black screen baseline. All results are thresholded at whole-brain family wise error corrected $p < 0.05$ . L: left. R: right. MOG: middle occipital gyrus. MTG: Middle temporal gyrus. P values are thresholded at $p < 0.05$ corrected for multiple comparisons using whole-brain family-wise error rate.

**Figure 2:**
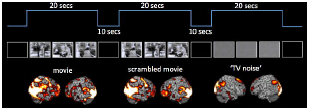
Block design results for movie, scrambled movie and ‘TV noise’ sequences. Figure displays overall brain activation for movie, scrambled movie and ‘TV noise’ sequences as measured in the block design paradigm. Results shown in a representative subject for T contrasts comparing movie, scrambled movie and ‘TV noise’ to a black screen baseline, thresholded at whole brain FWE corrected p<0.05.

### fMRI differentiation analysis

Figure 1, Bottom Panel, displays the experimental paradigm used for the differentiation analysis. In this paradigm, a 4 min sequence of movie, scrambled movie or ‘TV noise’ is presented to the subjects, each sequence repeated 30 times across different scanning sessions. In all three conditions, we expected an initial activation similar to that observed in the block design paradigm against the black screen baseline. However, unlike the block design analysis, the differentiation analysis focuses on systematic time-locked increases or decreases in activity compared to each voxel’s mean activity during the 4 min sequence. This is done in two ways: i) by statistically comparing BOLD activity values at each time point with the BOLD activity for a black screen baseline; ii) by statistically comparing BOLD activity values at each time point with the BOLD activity mean for the overall session. For the movie sequence, we expected that high phenomenological differentiation (many different experiences) would go along with high neurophysiological differentiation over time and space: each voxel would show significant activations/deactivations time-locked to specific movie frames (neurophysiological differentiation over time), and different voxels would do so for different frames (neurophysiological differentiation over space). For the scrambled movie sequence, we expected intermediate levels of neurophysiological differentiation, corresponding to intermediate levels of phenomenological differentiation. For the TV noise sequence, we expected minimal systematic time-locked activations/deactivations, with low neurophysiological differentiation (a single, unchanging pattern of activation/deactivation) going along with low phenomenological differentiation (unchanging experience of ‘TV noise’). Spontaneous fluctuations in BOLD activity from scan to scan were expected, but they would not be time locked to specific TV noise frames, which cortical regions should treat as equivalent.

For the differentiation analyses, fMRI data were analyzed with SPM and FMRIB Software Library (www.fmrib.ox.ac.uk/fsl) softwares and with additional scripts (MB, SS) written in Matlab (MathWorks Natick, MA). The first 4 volumes of each 4 min scanning session were first removed from the data set (allowing for T1 signal equilibration). Preprocessing of functional scans was performed as above using SPM for spatial realignment, normalization to Montreal National Institute template and 8 mm full width at half maximum (FWHM) smoothing. For all sessions, we then used FSL to remove linear trends by high pass filtering the data above a 60 sec. cutoff, and each voxel was then centered on its own session mean. Since we observed some non-specific T2* signal instability in the first 31 scans of each 4 min. scanning session, we excluded these volumes from our analysis [2].

We then measured Lempel-Ziv complexity on the spatiotemporal pattern of significant activations/deactivations. In a first analysis, we computed Lempel Ziv complexity for activations/deactivations compared to a black screen baseline (similar to the baseline used in the block design). These black screen values were taken from the final volume at the end each ten sec black screen presentation used as a baseline in the block design, which were themselves centered on the within-subject mean value of the block design session.

In order to identify differential activation patterns in response to different movie frames, we used SPM to perform F-tests between each given volume number across sessions and always the same black screen baseline. Statistical maps were obtained before Lempel-Ziv computation, as in [4], in order to only consider the deterministic part of the signal (systematic stimulus-induced changes in brain signals). Each F-test was thresholded at whole brain FWE corrected p<0.05 (as displayed in Figure 3 for a representative subject). Whole session results were then summarized in a binary spatio-temporal matrix of activations/deactivations, with each row corresponding to one voxel, and each column to an fMRI volume number in time. Lempel-Ziv complexity was computed on these spatio-temporal binary activation/deactivation matrices for movie, scrambled movie, and ‘TV noise’ (Figure 3 top, Table 2). Lempel-Ziv computation used a hierarchical clustering approach and each individual value was then normalized by the within-subject maximum before group mean and standard error of the mean were plotted for all conditions (Figure 4). Further analyses were performed to identify separately differential activations (positive deviations from the black screen baseline) and deactivations (negative deviations from the black screen) for each volume number across sessions. Positive and negative T-test contrasts were computed for each volume number and thresholded at FWE error corrected p<0.05. Lempel-Ziv complexity was then computed on binarized spatio-temporal activation or deactivation matrices for the movie, scrambled movie, and ‘TV noise’ conditions (Table 2).

**Table 2.**
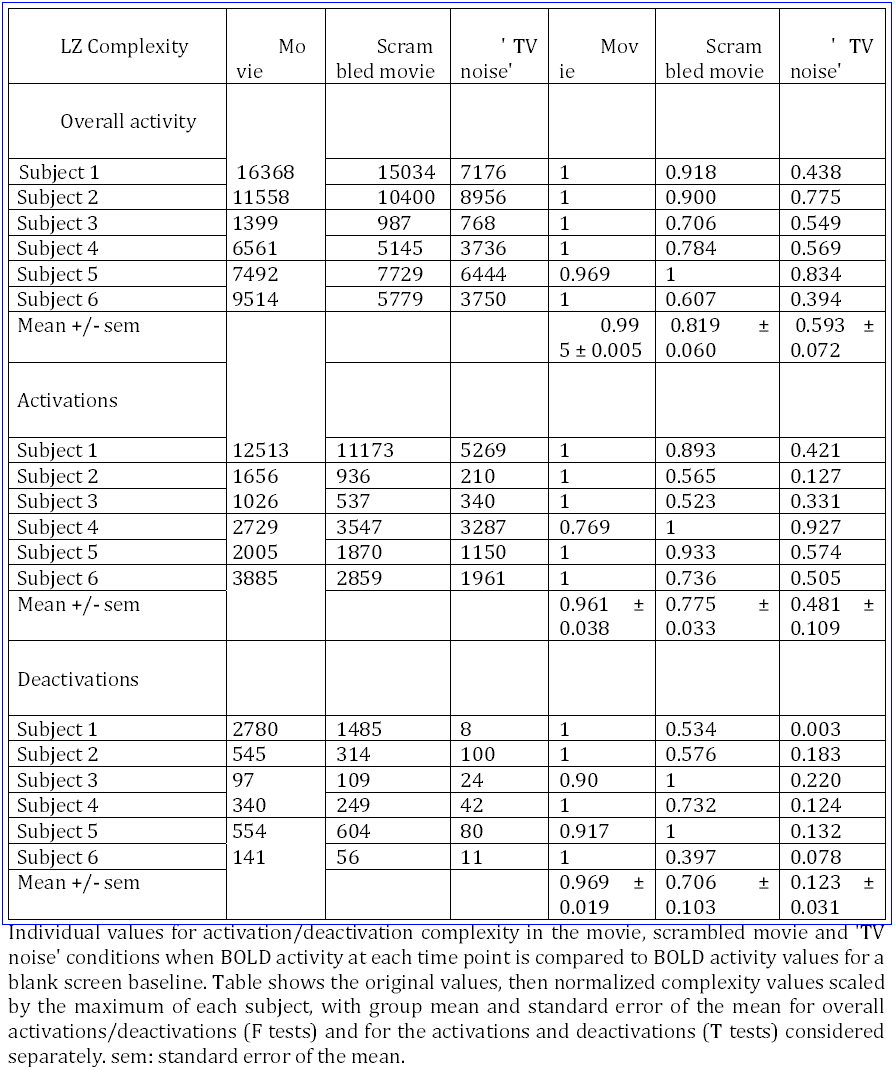
fMRI activation/deactivation Lempel-Ziv complexity results for the differentiation analysis comparing each volume BOLD signal to a black screen baseline.

**Figure 3.**
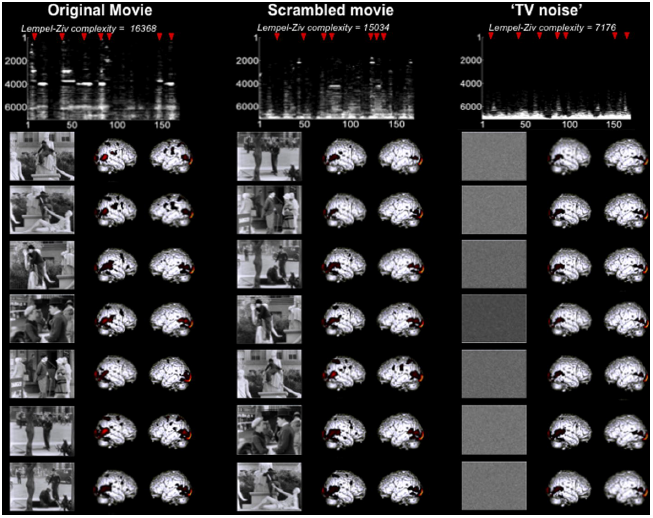
Lempel-Ziv complexity of brain activity correlates with stimulus set meaningfulness – comparison to a black screen baseline. Results shown for a representative subject (same subject as for Figure 2). Brain activity patterns over time are highly differentiated in the movie condition, intermediately differentiated in the scrambled movie condition, and very similar to one another in the ‘TV noise’ condition. Brain maps are here expressed in terms of significant changes in activity as compared to a black screen baseline (F-test, thresholded at whole brain FWE corrected p<0.05 for each frame). Top panel displays binarized spatio-temporal activation/deactivation matrices obtained for the 3 conditions after statistical thresholding was applied: a value of 1 was assigned to above threshold voxels for each scan, and a value of zero to voxels below threshold. For display purposes, binarized activation matrices are displayed only for the voxels that show at least once a significant activation in the movie (data dimension reduction from 94000 to ~7000 voxels). Lempel-Ziv complexity was computed at the whole brain activation matrix encompassing 94000 voxels in each condition.

**Figure 4.**
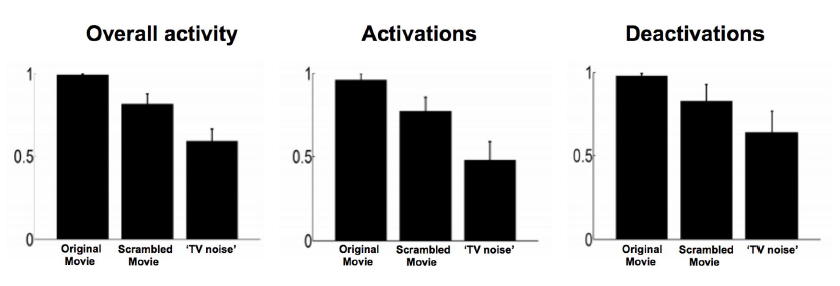
Lempel-Ziv complexity group values - comparison to a black screen baseline. Left panel: overall activations/deactivations (F test) group values. Middle panel: Lempel-Ziv complexity values for activations only (positive T test) Right panel: Lempel-Ziv complexity values for deactivations only (negative T test). For display purposes, each subject’s Lempel-Ziv complexity was normalized by its individual maximum value across all conditions. Bar graphs show group mean and standard error of the mean in each condition.

We then proceeded to a second, complementary differentiation analysis by performing, for each volume, an F-test of deviations from the session mean across the thirty sessions. In this analysis, significant non-zero values correspond to consistent positive or negative deviations in BOLD signal amplitude as compared to the session mean. As above statistical maps were obtained before Lempel-Ziv computation as in [4], in order to only consider the deterministic part of the signal. Again each F-test was thresholded at whole brain FWE corrected p<0.05 (as displayed in Figure 5) in order to perform a conservative correction for spatial multiple comparisons. Whole session results were then summarized in a binary spatio-temporal matrix of activations/deactivations. As in the first analysis, Lempel-Ziv complexity was computed on these spatio-temporal binary activation/deactivation matrices for movie, scrambled movie, and ‘TV noise’ (Figure 5 top, Table 3) using a hierarchical clustering approach. Each individual value was then normalized by the maximum before group mean and standard errors were computed for all conditions (Figure 6). Additional analyses were also performed to identify separately differential activations (positive deviations) and deactivations (negative deviations) as compared to the session mean for each volume number across sessions. Positive and negative T-test contrasts were computed for each volume number and as above thresholded at FWE error corrected p<0.05. Finally Lempel-Ziv complexity was computed separately on the binarized spatio-temporal activation or deactivation matrices for the movie, scrambled movie, and ‘TV noise’ conditions (see Table 3).

**Table 3.**
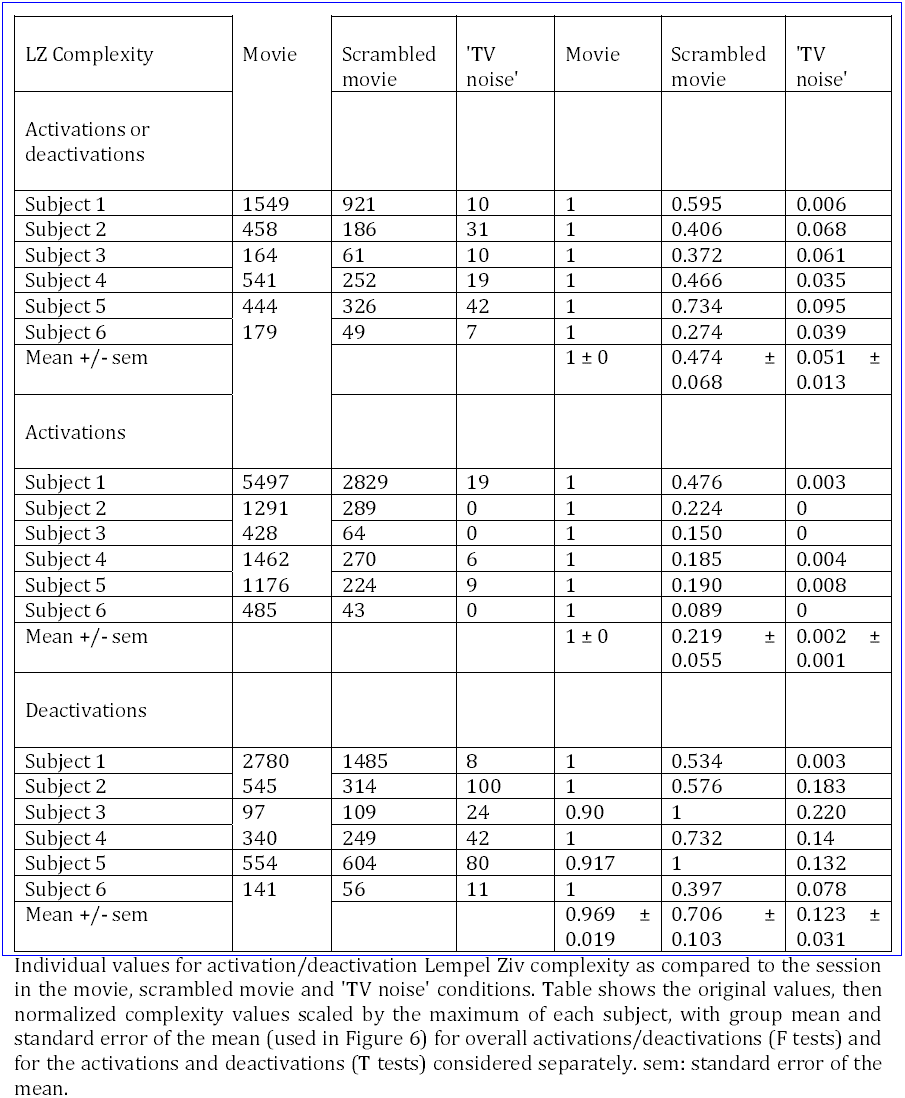
fMRI activation/deactivation Lempel-Ziv complexity results – comparison to the session mean

| LZ Complexity | Movie | Scrambled movie | 'TV noise' | Movie | Scrambled movie | 'TV noise' |
| --- | --- | --- | --- | --- | --- | --- |
| Activations or deactivations |  |  |  |  |  |  |
| Subject 1 | 1549 | 921 | 10 | 1 | 0.595 | 0.006 |
| Subject 2 | 458 | 186 | 31 | 1 | 0.406 | 0.068 |
| Subject 3 | 164 | 61 | 10 | 1 | 0.372 | 0.061 |
| Subject 4 | 541 | 252 | 19 | 1 | 0.466 | 0.035 |
| Subject 5 | 444 | 326 | 42 | 1 | 0.734 | 0.095 |
| Subject 6 | 179 | 49 | 7 | 1 | 0.274 | 0.039 |
| Mean +/- sem |  |  |  | 1 ± 0 | 0.474 ± 0.068 | 0.051 ± 0.013 |
| Activations |  |  |  |  |  |  |
| Subject 1 | 5497 | 2829 | 19 | 1 | 0.476 | 0.003 |
| Subject 2 | 1291 | 289 | 0 | 1 | 0.224 | 0 |
| Subject 3 | 428 | 64 | 0 | 1 | 0.150 | 0 |
| Subject 4 | 1462 | 270 | 6 | 1 | 0.185 | 0.004 |
| Subject 5 | 1176 | 224 | 9 | 1 | 0.190 | 0.008 |
| Subject 6 | 485 | 43 | 0 | 1 | 0.089 | 0 |
| Mean +/- sem |  |  |  | 1 ± 0 | 0.219 ± 0.055 | 0.002 ± 0.001 |
| Deactivations |  |  |  |  |  |  |
| Subject 1 | 2780 | 1485 | 8 | 1 | 0.534 | 0.003 |
| Subject 2 | 545 | 314 | 100 | 1 | 0.576 | 0.183 |
| Subject 3 | 97 | 109 | 24 | 0.90 | 1 | 0.220 |
| Subject 4 | 340 | 249 | 42 | 1 | 0.732 | 0.14 |
| Subject 5 | 554 | 604 | 80 | 0.917 | 1 | 0.132 |
| Subject 6 | 141 | 56 | 11 | 1 | 0.397 | 0.078 |
| Mean +/- sem |  |  |  | 0.969 ± 0.019 | 0.706 ± 0.103 | 0.123 ± 0.031 |

**Figure 5:**
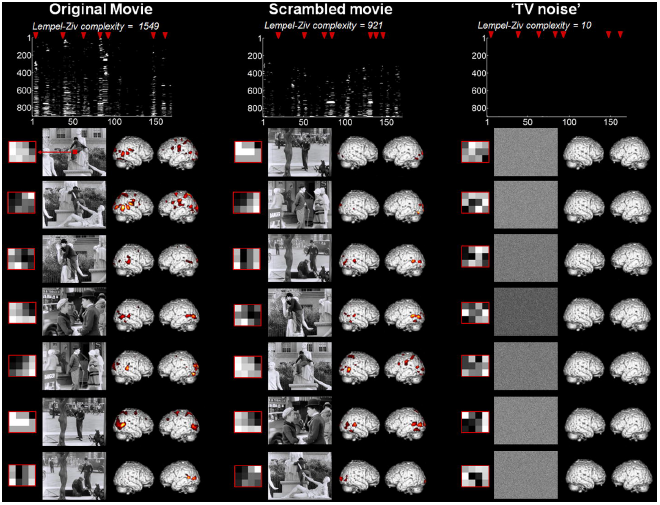
Lempel-Ziv complexity of brain activity correlates with stimulus set meaningfulness – comparison to the sequence mean. Figure displays differentiated brain activity patterns for movie, scrambled movie, and ‘TV noise’ stimulus sequences for the representative subject also used in Figure 2-3. Brain activity patterns over time are highly differentiated in the movie condition, intermediate in the scrambled movie condition, and very low in the ‘TV noise’ condition. Brain maps are here expressed in terms of significant changes in activity as compared to the within-session mean (F-test) thresholded at whole brain FWE corrected p<0.05 for each frame. Top panel displays binarized spatio-temporal activation/deactivation matrices obtained for the 3 conditions after statistical thresholding was applied – where a value of 1 was assigned to above threshold voxels for each scan, and a value of zero to voxels below threshold. For display purposes, binarized activation matrices are displayed only for the voxels that show at least once a significant activation in the movie (data dimension reduction from 94000 to ~900 voxels). Lempel-Ziv complexity was computed at the whole brain activation matrix encompassing 94000 voxels in each condition.

**Figure 6:**
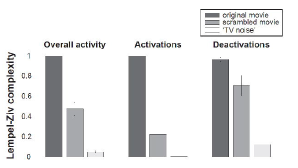
Group results for Lempel-ziv complexity analyses – comparison to the sequence mean. Lempel-Ziv complexity values for movie, scrambled movie, and ‘TV noise'. Left panel: overall activations/deactivations (F test) group values. Middle panel: complexity values for activations only (positive T test) Right panel: complexity values for deactivations only (negative T test). For display purposes, each subject’s Lempel-Ziv complexity was normalized by its individual maximum value across all conditions. Bar graphs show group mean and standard error of the mean in each condition.

### Lempel-Ziv complexity as a measure of differentiation

For the differentiation analyses, we calculated Lempel-Ziv complexity, a measure of the compressibility of a data set, for the spatiotemporal pattern of significant activations/deactivations with respect to a black screen baseline or to the session mean for the three conditions. Lempel-Ziv complexity was employed as a simple way to estimate the number of different activation/deactivation patterns.

Note that other measures, such as the total number of significantly activated/deactivated voxels, or the source entropy of the data, could also have been used to distinguish between the activation/deactivation patterns produced by our three conditions. In general, however, these other measures are not well suited to assess differentiation. For example, the number of activated voxels as well as source entropy, being only sensitive to first-order statistics, would be high for the voxel activation/deactivation patterns induced by a sequence alternating just two discriminable stimuli (say, a particular picture and black screen, repeated many times). By contrast, Lempel-Ziv complexity would immediately reveal the low differentiation (high compressibility) of the brain activation/deactivation patterns.

Note also that, in our previous work employing TMS-EEG to evaluate the differentiation of cortical EEG responses to transcranial magnetic stimulation [4], Lempel-Ziv complexity was normalized by source entropy. In that study, normalization was applied to control for variations in stimulation parameters (stimulation site and intensity) and in the behavioral state of the subjects (alert wakefulness, mild and deep anesthesia, sleep, and disorders of consciousness), so that, for the same amount of bran activation, we could estimate relative changes in complexity as reflecting the level of consciousness. In the present work, by contrast, the average intensity of the stimuli was similar across the three sequences, and the subjects’ level of consciousness was the same throughout the experiment. Moreover, our goal was to assess how many different patterns were triggered by the three stimulus sequences in absolute terms, rather than in relative terms. Specifically, we expected to observe many different patterns for the movie, hence high Lempel-Ziv complexity, a few less for the scrambled movie, and just one pattern throughout for the noise, hence low Lempel-Ziv complexity. For these reasons, we measured Lempel-Ziv complexity deliberately without normalization by source entropy.

Finally, to rule out that the hypothesized result - highest neurophysiological differentiation for the movie, intermediate for the scrambled movie, and lowest for ‘TV noise’ - could be accounted for simply by higher stimulus differentiation - we computed Lempel-Ziv complexity for the sequence of stimuli (images) in the three conditions. For computational expediency, we chose a subset of the pixels at the center of each image (one tenth of the full screen image size). Due to scrambling, we expected stimulus differentiation to behave opposite to neurophysiological differentiation, with the highest value for ‘TV noise’ and the lowest value for the movie.

### Integrated information Φ*

Integrated information is the amount of information generated by the whole above and beyond the information generated by its parts. A practical measure of integrated information, Φ*, can be defined as the difference between the mutual information of the whole system, I, and that of its parts, I* [5, 6],

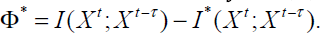

The first term, *I*(*X^t^*;*X^t-τ^*), quantifies how much information about the past state of a system can be decoded by knowing the present state (mismatch decoding). The second term, *I*^*^(*X^t^*;*X^t-τ^*, quantifies how much information about the past state can be decoded by knowing the present state, under the assumption that parts of the system are independent [7]. To do so, the past state of each part is decoded by using only its own present state, while ignoring the other parts of the system. If the parts are truly independent, I and I* are equal and integrated information, Φ*, is 0. If the parts interact with each other, there should be a difference between I and I*, and Φ* will be non zero. Φ* thus reflects how much information the system generates above and beyond its parts, i.e. its integrated information.

To calculate Φ*, fMRI time-courses of each voxel were averaged over 30 repetitions for movie, scrambled movie and ‘TV noise’. This was done in order to extract primarily the deterministic (stimulus-evoked) part of the time course of the responses, while discarding fMRI spontaneous activity fluctuations. Then, the mean time-courses of regions of interest (ROI) were calculated. We used two different sets of functionally relevant ROIs to confirm the robustness of our results. Each ROI was defined as a 5mm radius sphere around published coordinates. ‘ROI set 1’ included 123 ROIs which were defined based on fMRI resting state functional connectivity map [8]. ‘ROI set 2’ contained 160 ROIs that did not overlap with ROI set 1, which were identified by a meta-analysis of fMRI task-activation studies [9].

Quantifying causal interactions between regions across different time steps using fMRI is best performed while modeling regional differences in hemodynamic response function (HRF) [10, 11]. For this reason, we conducted a region-specific HRF deconvolution of our fMRI data before calculation of Φ* (using the approach described in [11], http://software.incf.org/software/blind-hrf-retrieval-and-deconvolution-for-resting-state-bold). This approach allows to deconvolve the data while taking into account regional differences in HRF shape, latency and FWHM and was designed in order to improve computation of effective connectivity on continuous signals using fMRI data [11].

To evaluate how much a system is integrated, one must find its informational ‘‘weakest link’’, i.e. the minimum information partition [MIP] [12], the partition of a system which makes the least of a difference. However, searching for the MIP exhaustively in large datasets is computationally infeasible. Furthermore, due to the small number of time points available in fMRI data, Φ* can only be computed on a limited number of ROIs. Given these limitations, we estimated Φ* by considering 1000 symmetric bipartitions of a set of 80 ROIs randomly selected from ROI set 1 or ROI set 2. For each bipartition, we calculated the amount of information generated by the set of 80 ROIs above and beyond its parts and that generated by the two sets of the 40 ROIs independently. The bipartition that made the least of a difference form the whole provided an approximate value of Φ*. We considered bipartitions only as they provide a lower bound on the expected value of integrated information [12]. We repeated this procedure for 1000 combinations of 80 ROIs from each ROI set in each condition, and computed the average Φ* values in each subject for each condition. Finally, we conducted group-level T tests in order to evaluate if average values of Φ* increased with changes in the meaningfulness of stimuli across conditions. Results were thresholded at false-discovery rate (FDR) corrected p<0.05 for each condition. This entire analysis was repeated five times through independent sampling of bipartitions to assess whether the results were robust.

### Neural Complexity

Neural Complexity *CN*(*X*) quantifies the average mutual information among bipartitions of a neural system [13]:

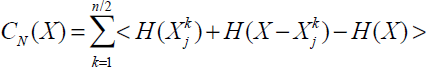

where *X* is the neural system with *n* elementary components, *Xjk* is a *j*-th subset consisting of *k* components, and *X-Xjk* is its complement in the system. *H(Xjk)*, *H(X-Xjk)* and *H(X)* are the entropies of *Xjk*, *X-Xjk* and *X* considered independently. <・> represents averaging for all subsets of size *k*. As done for Φ*, we calculated Neural Complexity on the deterministic (stimulus-evoked) part of fMRI time courses, i.e. the averaged time-courses over 30 repetitions for movie, scrambled movie and ‘TV noise’ conditions, both for ROI set 1 and ROI set 2. HRF deconvolution did not have to be performed to calculate Neural Complexity, since this measure is applied to instantaneous interactions between regions, with no delay in time (Neural Complexity results were nevertheless similar after deconvolution). Although theoretically Neural Complexity requires considering all bipartitions of the system, this computation is not practically feasible in fMRI data due to a combinatorial explosion. As we did for Φ*, we thus randomly selected bipartitions between two sets of 40 ROIs from ROI set 1 or ROI set 2 and calculated the mutual information between them. This procedure was repeated 10000 times to cover all ROIs of each ROI set. The 10000 values were averaged to obtain a single Neural Complexity estimate for each subject in each condition. Finally, we conducted group-level paired T tests in order to evaluate if Neural Complexity increased with the meaningfulness of the set of stimuli across conditions. Mean and standard error of the mean for these measures are displayed in Figure 5 for the movie, scrambled movie, and ‘TV noise’ conditions. Results were thresholded at false-discovery rate (FDR) corrected p<0.05 for each condition. Again, this entire analysis was repeated five times through independent sampling of bipartitions to assess whether the results were robust. Nevertheless, it should be emphasized that, even if Φ* and neural complexity values may be robust under repeated analysis, they remain extremely undersampled approximations.

## Results

### Lempel-Ziv complexity of the stimulus sequence

As expected, Lempel Ziv values computed over a representative fraction of the image pixels of the stimulus sequences (a central square) were lower for the movie sequence (1356), intermediate for the scrambled movie (1773), and highest for the ‘TV noise’ (2976).

### Block design analysis

The six subjects underwent a block design alternating a 20 seconds sequence of the movie (Chaplin’s ‘City Lights’), a 4-seconds time scrambled version of this movie sequence, and corresponding ‘TV noise’ sequences (see Figure 1, Upper Panel). In each subject, the movie, scrambled movie and ‘TV noise’ sequences all recruited a number of different brain areas (the results from an exemplar subject are shown in Figure 2). As shown in Table 1 (random effects group analysis), both movie and scrambled movie recruited more areas than TV noise, but did not differ from each other.

### fMRI activation Lempel-Ziv complexity

We then assessed the relationship between the differentiation of brain responses and the meaningfulness of the stimulus set. The same subjects watched a 4 minute sequence of the movie; a 4 minute sequence of the time-scrambled movie; a 4 minute sequence of the corresponding ‘TV noise’. Each sequence of movie, scrambled movie, and ‘TV noise’ was repeated 30 times in counterbalanced order (see Methods and Figure 1, Bottom Panel). After fMRI preprocessing, we centered each voxel on its own mean. A first differentiation analysis identified differentiated activation/deactivation as compared to a black screen baseline and a second analysis assessed BOLD increases or decreases as compared to the session mean (see Methods).

Figure 3 presents the results of a first kind of differentiation analysis, which assessed systematic activations/deactivations with respect to a common black screen baseline for the same representative subject shown in Figure 2. Clearly, the movie sequence produced many different activation/deactivation patterns that varied over time and space (high neurophysiological differentiation). By contrast, the TV noise sequence produced a pattern of activations/deactivations that was similar throughout the 4 min (low differentiation). The scrambled movie produced patterns that were less differentiated than the movie and much more differentiated that n TV noise.

We summarized the significant differential activation/deactivation patterns for each subject (F test) into a binarized spatio-temporal matrix (voxels x scans, where each voxel value is 1 if significant at family wise error (FWE) corrected p<0.05, and 0 if not significant for any given scan) for the three sets of stimuli (movie, scrambled movie and TV noise). Finally, we evaluated the Lempel-Ziv complexity of these binarized matrices. We repeated the process separately for activations (positive T test) and deactivations (negative T test) with respect to the black screen baseline. Figure 4 and Table 2 show the group results for this analysis. Highest group values of Lempel-Ziv complexity were found for the movie condition, intermediate values for the scrambled movie and lowest values for the ‘TV noise’ condition. Note that Lempel-Ziv complexity values for TV noise are much lower than for the other conditions, but they are not at zero. This is likely due to signal fluctuations, related to spontaneous activity and/or physiological noise.

A second kind differentiation analysis assessed systematic BOLD signal increases/decreases across sessions with respect to the mean of the sequence within each session. Figure 5 shows the results of this approach in the same subject shown in Figure 2 and 3. As in the first differentiation analysis, the movie sequence elicited over time a number of differentiated patterns of time-locked activation/deactivation widely distributed over the cortical surface. The scrambled movie sequence elicited time-locked activation/deactivation patterns that were less widespread. Finally, ‘TV noise’ induced virtually no significant differentiation of activation/deactivation patterns over the mean of each sequence.

We again summarized the significant differential activation/deactivation patterns for each subject (F test) into a binarized spatio-temporal matrix for the three sets of stimuli (movie, scrambled movie and TV noise) and computed the Lempel-Ziv complexity of these binarized matrices. We also computed Lempel-Ziv complexity separately for activations (positive T test) and deactivations (negative T test) as compared to session mean. Figure 5 (top) shows the results for this approach for overall activations/deactivations in our exemplar subject. Table 3 displays Lempel-Ziv complexity values for individual subjects. Figure 6 shows group means and standard errors. The results demonstrate that Lempel-Ziv complexity values of overall activations/deactivations, as well as activations and deactivations separately reflected the overall meaningfulness of the stimulus set, being highest for the original movie, intermediate for the scrambled movie, and low for TV noise.

### Integrated information Φ*

Neurophysiological activity patterns associated with conscious experiences should not only be differentiated, but also integrated, corresponding to a high capacity for information integration [4, 14, 15]. We estimated integrated information from fMRI data using a measure based on mismatch decoding, Φ* [5, 6]. Φ* is the difference between the information a system has about its past when taken as a whole and the information its parts have about themselves taken separately, considering the partition of the system that makes the least difference (minimum information partition, MIP). In other words, Φ* quantifies integrated information as the information the system has above and beyond its parts [14]. We chose two representative sets of 80 voxels (based on functional connectivity data (region of interest (ROI) set 1) and meta-analyses of activation (ROI set 2) respectively, see Methods) and performed two independent analyses. To calculate Φ*, we took the time series of mean BOLD values for each voxel averaged over the 30 repetitions for movie, scrambled movie, and TV noise. Total system information was defined as the mutual information between the state (BOLD signal values) of a set of ROI at each time step, and its state at an earlier time step. We repeated the analysis for intervals ranging from 1 to 10 seconds. The information for the parts independently was calculated in the same way. Integrated information Φ* was then defined as the difference between the total system information and the information for the minimum information partition (the partition, out of 1000 bipartitions sampled, which led to the least loss of information compared to the system as a whole [5, 6]. We used bipartition to calculate Φ*, because bipartition of the 80 ROIs provides a lower bound on the expected value of integrated information than any other partitions [12]. The results show that Φ* was highest for the movie, intermediate for the scrambled movie, and low for TV noise, for all time-lags up to 10 seconds (corrected P < 0.05, group data shown in Figure 7A for a time lag of 4 seconds, see also Table 4; for display purposes, results are normalized to the maximum Φ* value within each subject). Data for our exemplar subject at all time-lags are shown in Figure 7B. We obtained consistent results both for ROI set 1 and 2, which did not include any overlapping ROIs. Finally, to assess the robustness of the results, we repeated the entire analysis five times on independently chosen sets of bipartitions. In all cases, the results were confirmed: Φ* values for the movie condition were statistically higher than for scrambled, and both movie and scrambled were statistically higher than noise (corrected *P* < 0.05).

**Figure 7.**
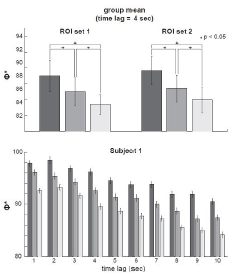
Group results for Integrated information Φ* analyses. Integrated information Φ* results for the movie, scrambled movie, and ‘TV noise’ conditions. Colors of bars represent conditions (dark gray: movie, medium gray: scrambled movie, light gray: TV noise). Upper left panel: the group-mean of Φ* calculated by using time lag of 4 second and ROI set 1. Error bar represents standard error of the mean for each measure in each condition. Asterisks indicate significant differences of the group means (p<0.05, corrected). Upper right panel: the group-mean of Φ* calculated by using time lag of 4 second and ROI set 2. Lower panel: Φ* calculated in our representative subject with ROI set 1, showing robust results across different time lags (1-10 seconds). In this panel, error bar indicates standard deviation of the mean for 1000 sets of 80 ROIs (see Methods).

**Table 4.** Integrated information Φ* results

| $\Phi^*$ | Movie | Scrambled movie | 'TV noise' | Movie | Scrambled movie | 'TV noise' |
| --- | --- | --- | --- | --- | --- | --- |
| ROI set 1 |  |  |  |  |  |  |
| Subject 1 | 96.1 | 92.5 | 89.5 | 1.000 | 0.963 | 0.932 |
| Subject 2 | 90.1 | 88.4 | 86.0 | 1.000 | 0.981 | 0.955 |
| Subject 3 | 92.8 | 90.8 | 87.5 | 1.000 | 0.979 | 0.943 |
| Subject 4 | 83.9 | 81.8 | 81.4 | 1.000 | 0.975 | 0.970 |
| Subject 5 | 84.4 | 83.8 | 85.2 | 0.991 | 0.984 | 1.000 |
| Subject 6 | 86.6 | 84.8 | 82.6 | 1.000 | 0.979 | 0.954 |
| Mean +/- sem |  |  |  | 0.999±<br>0.002 | 0.977±<br>0.003 | 0.959±<br>0.010 |
| ROI set 2 |  |  |  |  |  |  |
| Subject 1 | 94.8 | 91.4 | 91.1 | 1.000 | 0.964 | 0.961 |
| Subject 2 | 93.1 | 92.0 | 88.9 | 1.000 | 0.989 | 0.955 |
| Subject 3 | 92.8 | 89.8 | 87.5 | 1.000 | 0.968 | 0.943 |
| Subject 4 | 86.5 | 83.5 | 84.1 | 1.000 | 0.964 | 0.972 |
| Subject 5 | 84.7 | 83.8 | 83.7 | 1.000 | 0.989 | 0.988 |
| Subject 6 | 86.1 | 83.8 | 80.4 | 1.000 | 0.974 | 0.934 |
| Mean +/- sem |  |  |  | 1.000 | 0.975±<br>0.005 | 0.959±<br>0.008 |

### Neural Complexity

Finally, we asked whether Neural Complexity, another measure of information integration within a network [13], was also correlated with stimulus set meaningfulness, using the same datasets as for the calculation of Φ*. While Φ*considers the mutual information between current and past states of the whole system and its parts, Neural Complexity measures the average mutual information between the current states of one part of the system and the rest. As shown in Figure 8 and Table 5, the group mean for Neural Complexity was again high for the movie, intermediate for the scrambled movie, and low for ‘TV noise’ (for display purposes, results were again normalized to the maximum Neural Complexity value within each subject). Thus, the overall meaningfulness of the stimulus set was reflected in measures of information integration among cortical regions, such as Φ* and Neural Complexity, indicating that differentiated responses to the movie were also integrated. As above, to assess the robustness of the results, we repeated the entire analysis five times on independently chosen sets of bipartitions. In all cases, the results were confirmed: neural complexity values for the movie condition were statistically higher than for scrambled, and both movie and scrambled were statistically higher than noise (corrected P < 0.05).

**Figure 8:**
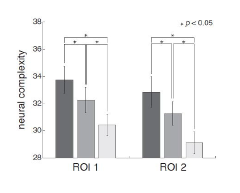
Group results for Neural Complexity analyses. Neural Complexity results for the movie, scrambled movie, and ‘TV noise’ conditions. Colors of bars represent conditions (dark gray: movie, medium gray: scrambled movie, light gray: TV noise). Left panel: group means and standard deviation of the mean for Neural Complexity calculated on ROI set 1. Right panel: group means and standard deviation of the mean for Neural Complexity calculated on ROI set 2. Asterisks indicate significant differences of the group means (p<0.05, corrected).

**Table 5.** Neural Complexity results

| Neural Complexity | Movie | Scrambled movie | 'TV noise' | Movie | Scrambled movie | 'TV noise' |
| --- | --- | --- | --- | --- | --- | --- |
| ROI set 1 |  |  |  |  |  |  |
| Subject 1 | 35.4 | 33.9 | 31.8 | 1.000 | 0.957 | 0.897 |
| Subject 2 | 36.7 | 34.0 | 31.8 | 1.000 | 0.928 | 0.866 |
| Subject 3 | 34.5 | 34.9 | 32.3 | 0.989 | 1.000 | 0.927 |
| Subject 4 | 31.9 | 31.0 | 30.4 | 1.000 | 0.972 | 0.954 |
| Subject 5 | 29.9 | 29.4 | 27.7 | 1.000 | 0.983 | 0.927 |
| Subject 6 | 34.2 | 30.4 | 28.7 | 1.000 | 0.890 | 0.839 |
| Mean +/- sem |  |  |  | 0.998±<br>0.002 | 0.955±<br>0.016 | 0.902±<br>0.018 |
| ROI set 2 |  |  |  |  |  |  |
| Subject 1 | 34.7 | 32.6 | 29.8 | 1.000 | 0.940 | 0.860 |
| Subject 2 | 35.8 | 32.9 | 30.5 | 1.000 | 0.919 | 0.853 |
| Subject 3 | 33.8 | 33.7 | 31.4 | 1.000 | 0.999 | 0.929 |
| Subject 4 | 32.3 | 30.4 | 29.3 | 1.000 | 0.942 | 0.908 |
| Subject 5 | 28.0 | 28.7 | 26.3 | 0.974 | 1.000 | 0.916 |
| Subject 6 | 32.6 | 29.2 | 27.3 | 1.000 | 0.897 | 0.838 |
| Mean +/- sem |  |  |  | 0.996±<br>0.004 | 0.950±<br>0.017 | 0.884±<br>0.016 |

## Discussion

This study tested the hypothesis that the differentiation of cortical responses to a set of diverse stimuli reflects the meaningfulness of the stimulus set for the subject. Objectively, the three stimulus sets – movie, scrambled movie, and ‘TV noise’ – consisted in each case of many different frames. In fact, the differentiation of the stimulus set, as measured by Lempel-Ziv complexity, was lowest for the movie and highest for TV noise, as one would expect due to stimulus scrambling. From the subject’s perspective, however, while the movie sequence consisted of a set of different ‘scenes’ (phenomenological differentiation), the ‘TV noise’ sequence consisted of just a single experience, that of seeing ‘TV noise’ (lack of phenomenological differentiation). Thus the movie sequence was rich in diverse meanings, the ‘TV noise’ had just one meaning, and the scrambled movie was in between. Our findings demonstrate that the differentiation of cortical activity patterns was indeed highest for a sequence of frames from a movie, intermediate for a temporally scrambled sequence of the same movie, and minimal for a ‘TV noise’ sequence obtained by spatially scrambling the pixels from each movie frame – the opposite ranking compared to stimulus differentiation. Our results support the hypothesis that neurophysiological differentiation reflects phenomenological differentiation and the overall meaningfulness of a set of stimuli.

All three sets of stimuli induced reproducible activation of multiple brain regions compared to a black screen baseline (block design analysis, Fig. 2). Nevertheless, the differentiation of brain activations, measured using Lempel-Ziv complexity, was highest for movie, intermediate for scrambled, and minimal for ‘TV noise’ (Fig. 3-6). Lempel-Ziv complexity assesses the compressibility of data – here, fMRI cortical activation patterns. We employed this measure to evaluate the absolute diversity of activation/deactivation patterns because, unlike some other measures such as overall activation or source entropy, it is sensitive to more than first-order statistics (see Methods). In general, a high Lempel-Ziv value (as obtained for the set of movie frames) indicates that different stimuli in the set induced many different activation/deactivation patterns, which are hard to compress. At the other extreme, the set of ‘TV noise’ stimuli, while just as different from each other as movie frames, induced a cortical activation/deactivation pattern that did not change significantly from one frame to the next, and was thus easy to compress. It should be noted that changes in higher order image statistics due to spatial scrambling in the ‘TV noise’ condition may contribute to some of the observed differences (see methods). From a theoretical perspective, however, any change in neurophysiological differentiation due to the scrambling of regularities – including low-level ones – reflect stimulus meaningfulness [16]. At any rate, the effects of changes in low-level spatial statistics appear to be confined to early visual areas [17], which contribute only a fraction of the voxels evaluated in our analysis. Also, the increased neurophysiological differentiation in the movie compared to the time-scrambled movie cannot be accounted for by changes in spatial statistics.

Much evidence indicates that a characteristic set of regions - the default mode network - is deactivated in several different task conditions [18]. Here we also observed a higher differentiation of cortical responses to the movie compared to ‘TV noise’ not just for activations but also for deactivations. This implies that different meaningful stimuli turn off different sets of brain areas. By contrast, a set of stimuli having the same meaning (‘TV noise’) turns off the same set of areas. Therefore, in addition to the task-related deactivation of a default brain network, there can be spatially differentiated, stimulus-specific deactivations that relate to stimulus meaning.

The higher Lempel-Ziv complexity of movie over ‘TV noise’ was paralleled by a higher value of integrated information Φ* among cortical regions (Fig. 7, [5, 6]). Φ* is a measure of how much better the current state of a system as a whole predicts its future state compared to what independent parts of the system could do, for the partition that cuts the system through its weakest link (minimum information partition, MIP). For Φ* to be high, a system must have a large repertoire of different states (information) and any part of a system must interact effectively with the rest of the system (integration). Thus, like Lempel-Ziv complexity, Φ* reflects the degree of differentiation of brain responses, while at the same time establishing that such responses are integrated. This finding suggests that the higher differentiation observed in the movie condition is due to the coherent rather than to the independent involvement of distributed brain areas. Another analysis consistent with this interpretation was the higher average mutual information between a part of the cortex and the rest in the movie compared to scrambled movie and ‘TV noise’ (Neural Complexity, Fig. 8). As shown in theoretical work and computer simulations [19], increases in mutual information between cortical regions when the brain is exposed to meaningful stimuli are proportional to changes in the average mutual information between the stimulus set and subsets of cortical regions. This is true even though, by the data processing theorem, the overall mutual information between the stimulus set and the brain is fixed, and is comparable in the three experimental conditions [20]. Thus, an increase in average mutual information across partitions of the cortex in the movie condition indicates that the information in the stimulus set must be distributed more efficiently to different brain regions, where each region deals with selective aspects of the same stimuli [19]. This fits with the intuitive idea that different meaningful stimuli should be informative in different ways for different brain regions [21].

We recently employed Lempel-Ziv complexity to evaluate the differentiation of cortical responses after transcranial magnetic stimulation (TMS) activation of individual brain regions [4]. Compared to measures of functional connectivity, Lempel-Ziv complexity has the advantage of combining in a single measure the spread of activity and the differentiation of brain responses to perturbations [4], both of which are necessary for information integration [22]. The results showed that Lempel-Ziv complexity is high when subjects are conscious and low when consciousness is lost. This was true for individual subjects and across different conditions such as loss of consciousness with sleep, general anesthetics, and brain damage. In the TMS work, a direct cortical perturbation, which in itself evokes no conscious content, was employed to gauge the level of consciousness, as reflected in the brain’s capacity for integration and differentiation [4]. In the current study, the differentiation of brain responses was evaluated in conscious subjects in response to different sensory stimuli, each of which evoked a conscious content, to assess the meaningfulness of a set of stimuli for the subject.

The assessment of the differentiation of cortical responses to stimuli departs from standard fMRI localization approaches, since it does not ask which particular brain areas are activated by particular features or categories contained in different stimuli. Also, unlike decoding approaches, our analysis does not ask whether it is possible to infer the nature of the stimuli from brain activity. Nevertheless, for this study we chose stimuli such that the differences in phenomenological differentiation/meaningfulness were obvious (movie > scrambled movie > TV noise) and similar across different subjects. On the other hand, some of the meaningful categories present in the movie and absent in ‘TV noise’ were also obvious (e.g. faces and houses). We also expected that different movie frames would activate different cortical regions at different times, such as the fusiform face area (FFA) for frames with faces and the parahippocampal place area (PPA) for frames with houses, whereas ‘TV noise’ frames would not activate these areas differentially. Finally, based on the pioneering work by Hasson et al. [2, 23, 24], we expected that high level areas characterized by long temporal receptive fields would be differentially activated between all conditions. These expectations could have been confirmed through a traditional localization or decoding approach, but our purpose here was different: to exploit such differential responses to validate the notion that the differentiation of neurophysiological responses can be measured and that it reflects phenomenological differentiation, hence meaningfulness. In this respect, it is instructive to compare the movie and the scrambled movie condition. In the block design, there was no significant difference in regional activation/deactivation between the two (Table 1). Moreover, it is difficult to say exactly what categories of meaning were missing in the scrambled sequence and where each of them would map on the cortex. However, neurophysiological differentiation clearly distinguished between the two sets according to their meaningfulness. More generally, assessing the differentiation of brain responses should be helpful when it is unclear a priori which stimuli might be ecologically relevant and how they may be categorized by the brain, for example in infants or animals. It can also be useful when brain responses are highly variable across individuals or idiosyncratic, as when the meaning of stimuli is strictly personal, or when brain injuries lead to alterations in cortical organization. In such cases, while localization of responses or decoding what the subjects may be perceiving may be difficult, it may be possible to establish whether some sets of stimuli are more or less meaningful than others. Finally, this approach should be especially powerful when comparing the differentiation of the neurophysiological responses of different subjects to the same stimuli, indicating which stimulus set may be more meaningful to which subject. As in this study, ‘TV noise’ can provide a common baseline representing a bound of least meaningfulness. It should be emphasized that the indices employed here to quantify the differentiation of neurophysiological responses are meant to capture the overall meaningfulness of a set of stimuli for a subject, not the actual meaning of a particular stimulus. The relationship between the quantity or level of meaningfulness and the quality of a particular meaning is analogous to that between the quantity or level of consciousness and the quality of a particular experience, which can only be specified by characterizing the associated conceptual structure [16].

In conclusion, these results suggest that the differentiation of neural responses can reflect the meaningfulness of a given set of stimuli for a given subject, without prior assumptions about which stimuli might be relevant and how and where they may be represented in the brain. Forthcoming experiments using both fMRI and high-density EEG will assess neurophysiological differentiation using a larger spectrum of stimulus sets, including sets that may have different levels of meaningfulness for different subjects. Future studies will also take advantage of single trial presentation of stimuli rather than repeated ones, and employ state space analyses (similar to the representational analysis used in fMRI decoding [25]) to quantify differentiation in a multivariate manner. If this approach proves successful, it will become feasible to employ specifically designed sets of stimuli to explore what features of the environment may be most meaningful for a given subject, and compare the maximum amount of differentiation and therefore meaning that different subjects can extract from different sets of stimuli. Finally, similar experimental paradigms could be used to investigate in which set of brain areas neurophysiological and phenomenological differentiation covary most closely, providing a way to locate the neural substrates of consciousness and subjective meaning without having to rely on explicit report.

## Acknowledgments

Supported by a Mind Science Foundation grant to MB and GT, a Postdoctoral Fellowship from FNRS and a University of Liege travel grant to MB, and by a McDonnell Foundation grant to GT. OG received support from NIH grant MH095984 to Bradley R. Postle and Giulio Tononi, Belgian National Funds for Scientific Research (FNRS) and Fonds Léon Fredericq. Grants-in-Aid for the Japan Society for the Promotion of Science (JSPS) Fellows 2311065 and a Nakayama Foundation for Human Science grant to SS. The authors declare no conflict of interest.

## References

1. Tononi G (2012) The integrated information theory of consciousness: an updated account. Arch Ital Biol 150: 56–90.

2. Hasson U, Yang E, Vallines I, Heeger DJ, Rubin N (2008) A hierarchy of temporal receptive windows in human cortex. J Neurosci 28: 2539–2550.

3. Groen, II, Ghebreab S, Lamme VA, Scholte HS (2012) Spatially pooled contrast responses predict neural and perceptual similarity of naturalistic image categories. PLoS Comput Biol8: e1002726.

4. Casali AG, Gosseries O, Rosanova M, Boly M, Sarasso S, et al. (2013) A theoretically based index of consciousness independent of sensory processing and behavior. Sci Transl Med 5: 198ra105.

5. Oizumi M, Ishii T, Ishibashi K, Hosoya T, Okada M (2010) Mismatched decoding in the brain. J Neurosci 30: 4815–4826.

6. Oizumi M, Yanagawa T, Amari S, Tsuchiya N, Fujii N (2012) Measuring the level of consciousness based on the amount of integrated information computed from electrocorticogram (ECoG) recordings in monkeys before and after anesthesia. Neuroscience New Orleans, USA.

7. Oizumi M, Okada M, Amari S (2011) Information loss associated with imperfect observation and mismatched decoding. Front Comput Neurosci 5: 9.

8. Dosenbach NU, Nardos B, Cohen AL, Fair DA, Power JD, et al. (2010) Prediction of individual brain maturity using fMRI. Science 329: 1358–1361.

9. Power JD, Cohen AL, Nelson SM, Wig GS, Barnes KA, et al. (2011) Functional network organization of the human brain. Neuron 72: 665–678.

10. David O, Guillemain I, Saillet S, Reyt S, Deransart C, et al. (2008) Identifying neural drivers with functional MRI: an electrophysiological validation. PLoS Biol 6: 2683–2697.

11. Wu GR, Liao W, Stramaglia S, Ding JR, Chen H, et al. (2013) A blind deconvolution approach to recover effective connectivity brain networks from resting state fMRI data. Med Image Anal 17: 365–374.

12. Balduzzi D, Tononi G (2008) Integrated information in discrete dynamical systems: motivation and theoretical framework. PLoS Comput Biol 4: e1000091.

13. Tononi G, Sporns O, Edelman GM (1994) A measure for brain complexity: relating functional segregation and integration in the nervous system. Proc Natl Acad Sci U S A 91: 5033–5037.

14. Tononi G (2008) Consciousness as integrated information: a provisional manifesto. Biol Bull 215: 216–242.

15. Barrett AB, Seth AK (2011) Practical measures of integrated information for time-series data. PLoS Comput Biol 7: e1001052.

16. Tononi G (2012) Integrated information theory of consciousness: an updated account. Arch Ital Biol 150: 293–329.

17. Musel B, Bordier C, Dojat M, Pichat C, Chokron S, et al. (2013) Retinotopic and lateralized processing of spatial frequencies in human visual cortex during scene categorization. J Cogn Neurosci 25: 1315–1331.

18. Buckner RL, Andrews-Hanna JR, Schacter DL (2008) The brain’s default network: anatomy, function, and relevance to disease. Ann N Y Acad Sci 1124: 1–38.

19. Tononi G, Sporns O, Edelman GM (1996) A complexity measure for selective matching of signals by the brain. Proc Natl Acad Sci U S A 93: 3422–3427.

20. Cover T, Thomas J (1991) Elements of Information Theory; Wiley, editor: Wiley.

21. Friston KJ, Tononi G, Sporns O, Edelman GM (1995) Characterising the complexity of neuronal interactions. Human Brain Mapping 3: 302–314.

22. Oizumi M, Albantakis L, Tononi G (2014) From the phenomenology to the mechanisms of consciousness: integrated information theory 3.0. PLoS Comput Biol 10: e1003588.

23. Hasson U, Malach R, Heeger DJ (2010) Reliability of cortical activity during natural stimulation. Trends Cogn Sci 14: 40–48.

24. Honey CJ, Thesen T, Donner TH, Silbert LJ, Carlson CE, et al. (2012) Slow cortical dynamics and the accumulation of information over long timescales. Neuron 76: 423–434.

25. Haxby JV, Guntupalli JS, Connolly AC, Halchenko YO, Conroy BR, et al. (2011) A common, high-dimensional model of the representational space in human ventral temporal cortex. Neuron 72: 404–416.

